# Robust survival-based RNAi using in tandem silencing of adenine phosphoribosyltransferase

**DOI:** 10.1101/2020.04.09.034132

**Authors:** Robert G. Orr, Stephen J. Foley, Giulia Galotto, Boyuan Liu, Luis Vidali

**Author notes:** Corresponding author information: Luis Vidali, Associate Professor in Department of Biology and Biotechnology, Worcester Polytechnic Institute, 60 Prescott Street, Worcester, MA 01609. Author contributions: R.G.O., and L.V. designed and supervised the research. L.V. conceived the project. S.J.F. created the APTi constructs. R.G.O. developed and optimized the APTi microscope assay and western blots. G.G., and B.L. contributed to initial optimization of microscope assay. R.G.O wrote the manuscript with contributions from S.J.F., G.G., and L.V.

## Abstract

RNA interference (RNAi) enables flexible and dynamic interrogation of entire gene families or essential genes without the need for exogenous proteins, unlike CRISPR-Cas technology. Unfortunately, isolation of plants undergoing potent gene silencing requires laborious design, visual screening, and physical separation for downstream characterization. Here, we developed a novel APT-based RNAi technology (APTi) in *Physcomitrella patens* that simultaneously improves upon the multiple limitations of current RNAi techniques. APTi exploits the pro-survival output of transiently silencing the APT gene in the presence of 2-fluoradenine, thereby establishing survival itself as a reporter of RNAi. To maximize silencing efficacy of gene targets we created vectors that facilitate insertion of any gene target sequence in tandem with the APT silencing motif. The APTi approach resulted in a homogenous population of *P. patens* mutants specific for our gene target, with zero surviving background plants within 8 days. The observed mutants directly corresponded to a maximal 93% reduction of the tested target protein, substantially exceeding previous dsRNA methods. The positive selection nature of APTi represents a fundamental improvement in RNAi technology and will contribute to the growing demand for technologies amenable to high-throughput phenotyping.

**One-sentence summary:** Generation of dsRNA targeting the *APT* gene in tandem with a target gene enables positive selection of strongly silencing plants.

## Introduction

Loss-of-function studies have long served as building blocks of our understanding of biological processes. Currently, the researcher is confronted with an ever-growing toolbox to perturb gene function at the genetic, transcript, or protein level, with each presenting its unique challenges and insights (Housden et al., 2017). CRISPR-Cas technology has facilitated targeted genetic knockouts in previously intractable systems, but its efficacy is limited by accessibility of the genetic locus and is subject to off-target effects (Horlbeck et al., 2016; Jensen et al., 2017; Verkuijl and Rots, 2019). Furthermore, genetic knockouts cannot isolate essential genes or reversibly interrogate developmental, tissue, or cell specific functions of a given gene or splice variant. Importantly, recent work has questioned the lasting dogma that genetic knockouts are the “gold standard” loss-of-function approach (Rossi et al., 2015; Smits et al., 2019). These studies demonstrated genetic compensation or residual protein expression, signifying that genetic alterations alone are insufficient to directly infer gene function. Therefore, reliable inferences of loss-of-function studies will require integration of multiple independent approaches (Deans et al., 2016), ideally in a high-throughput manner to maximize confidence.

RNA interference (RNAi) is a popular gene silencing strategy that does not depend upon exogenous proteins, such as Cas9, and enables reversible reduction of protein levels through targeted degradation of mRNA (Small, 2007). RNAi’s ease of use and flexibility lends itself as an indispensable complement to genetic editing techniques. Traditionally, RNAi in plants is induced through expression of long inverted repeats that self-base pair to form double-stranded RNA (dsRNA), which is then processed into multiple small interfering RNAs (siRNA) and targeted to complementary sequences within mRNA (Chuang and Meyerowitz, 2000; Hannon, 2002; Baulcombe, 2004). Importantly, a single dsRNA targeting one gene can simultaneously silence multiple genes with sufficient similarity, or a single dsRNA can be generated that contains multiple gene targets in tandem for simultaneous silencing (Vidali et al., 2007; Li et al., 2013). The expression of dsRNAs can be specifically modulated, either through induction or unique promoters, which allows developmental and cell-type specific reduction of even essential gene products (Byzova et al., 2004; Nakaoka et al., 2012; Miki et al., 2015; Liu and Yoder, 2016). Despite these advantages, RNAi is hindered by variable efficacy of target gene silencing and potential off-target effects (Xu et al., 2006). However, some argue the prevalence of off-target effects is overstated (Zimmer et al., 2019), and importantly an ideal RNAi experiment should demonstrate rescue of the gene silencing phenotype (Vidali et al., 2009; Vidali et al., 2010; Ding et al., 2018). Work using artificial microRNAs (amiRNA) attempts to circumvent the limitations of traditional dsRNA-based RNAi by engineering a single siRNA (Schwab et al., 2006; Gutierrez-Nava et al., 2008). Although amiRNA technology ameliorates possible off-targets derived from the initial dsRNA, evidence in *Physcomitrella patens* showed generation of additional siRNAs upon cleavage of the amiRNA target, potentially negating the specificity of the amiRNA (Khraiwesh et al., 2008). Furthermore, amiRNAs display variable silencing efficiency, thereby necessitating screening of multiple amiRNAs and limiting experimental throughput (Li et al., 2013; Zhang et al., 2018). To date, no RNAi method addresses another potential source of variability of silencing: transcriptional silencing of the RNAi transgene mediated by DCL3 (Morel et al., 2000; Fusaro et al., 2006; Small, 2007).

Generation and characterization of loss-of-function mutants using gene silencing methods is fundamentally a two-step process: (1) the molecular mechanism resulting in gene silencing; (2) the identification and isolation of your target undergoing gene silencing to further characterize. Recent advancements in gene silencing technologies focused on enhancing the flexibility and robustness of (1) (Hauser et al., 2013; Zhang et al., 2018) but fail to address the practical limitations imposed by (2). Identification of actively silencing mutants has been eased by coupling the silencing target of interest to a reporter, such as a nuclear-localized fluorescent protein (Bezanilla et al., 2005; Vidali et al., 2007; Vidali et al., 2010; van Gisbergen et al., 2018; Zhang et al., 2018). When paired with an automated or semi-automated image acquisition and analysis pipeline the burden of identifying silencing mutants is substantially mitigated (Wu and Bezanilla, 2012; Galotto et al., 2019). Nevertheless, the tedium of manually isolating silencing plants remains. This limitation is a consequence of traditional gene-silencing construct design, whereby the silencing module is regulated independently from the selectable marker. A typical gene-silencing experiment will contain a heterozygous population of actively silencing plants, presumably a result of the plant silencing the exogenous silencing module to rescue itself (Fusaro et al., 2006; Khraiwesh et al., 2010). This transcriptional-based silencing not only increases variability of target silencing, but when coupled with visual screening it exacerbates the manual labor required to isolate mutants. Therefore, downstream characterization of silencing plants, such as qPCR and western blots is stymied. Furthermore, certain reporter-based silencing limits the experimental scope to testing only within established reporter lines, (Bezanilla et al., 2003; Nakaoka et al., 2012). Finally, using a fluorescent reporter to infer silencing could complicate the subsequent use of fluorescent reporters, such as biosensors or fluorescent fusions for protein localization, to characterize mutant function.

Here, we generated a modular, Gateway-based, RNAi construct that couples silencing any gene(s) of interest in tandem with silencing of adenine phosphoribosyltransferase (APT-interference/APTi). This approach results in near undetectable levels of background non-silencing transformants, thereby trivializing the identification and isolation of actively silencing plants to the simple observation and harvesting of all living plants. Unlike traditional antibiotic positive selection followed by visual screening, our new construct simultaneously selects for transformation and silencing through positive selection alone. We achieved this by exploiting the ubiquitous selectable marker system APT, which converts purine analogs to cytotoxic nucleotides (Schaff, 1994). The APT loss-of-function selection system has been successfully used in plants (Moffatt and Somerville, 1988; Charlot et al., 2014), mammals (Schaff et al., 1990), and bacteria (Levine and Taylor, 1982), but to our knowledge all experiments with APT involve stable genetic mutants. To quantitatively evaluate the performance of our APTi system, we used the plant model organism *Physcomitrella patens* and its dramatic *myosin XI* phenotype (Vidali et al., 2010). We show that transiently silencing endogenous APT is sufficient for efficient positive selection and facilitates downstream characterization of mutants. Our APTi system resulted in >90% reduction of target myosin XI protein abundance, which far surpasses other dsRNA-based methods (Vidali et al., 2007; Vidali et al., 2010; Nakaoka et al., 2012; Guo et al., 2019) and is comparable to silencing efficiencies for optimized amiRNAs (Zhang et al., 2018).

## Results

### Development of A Novel Positive Selection RNAi Methodology Based on Endogenous APT Activity

Current RNAi methods produce a range of phenotype severity due to variable silencing efficiencies. This variability necessitates optimization experiments that screen fluorescent reporters to maximize silencing by gene sequence targets (Li et al., 2013; Zhang et al., 2018). We sought to simultaneously improve RNAi silencing efficiency and streamline characterization of RNAi mutants by coupling silencing of the gene target with a survival advantage. Previous work in *P. patens* directly coupled the sequence of a stably integrated, constitutively expressed reporter, such as a fluorescent protein and/or GUS, to the gene target sequence in inverted repeats (Bezanilla et al., 2003; Bezanilla et al., 2005; Nakaoka et al., 2012). Therefore, expression would result in dsRNA formation and co-reduction of the intracellular reporter and the coupled target gene. We predicted that coupling the sequence of a lethal reporter to any gene target would result in maximal co-silencing to ensure silencing of the lethal reporter, thereby promoting survival.

The adenine phosphoribosyltransferase (APT) gene has been frequently used as a reporter to evaluate gene-targeting efficiency (Schaefer, 2001; Charlot et al., 2014). Functional APT converts adenine analogs, such as 2-fluoroadenine (2-FA), to cytotoxic nucleotides (Schaff, 1994). Therefore, sufficient reduction of APT activity will impart resistance to 2-FA, but this has only been demonstrated in genetic knockouts. To test if silencing of APT effectively conferred survival to plants grown on 2-FA, we inserted an APT targeting sequence into a previously developed RNAi vector that also contains a reporter targeting sequence (Bezanilla et al., 2005). To maximize silencing efficacy, we generated an APT targeting sequence consisting of the 5’ UTR (179 bp) and first 210 bp of the APT gene. The APT silencing construct (APT-RNAi) conferred resistance to wild-type *P. patens* cultured on 1.25 μg/mL 2-FA, whereas zero plants survived on 2-FA when transformed with a control plasmid lacking the APT silencing sequence (Control-RNAi) (Fig. 1A). This result clearly establishes survival on 2-FA paired with APT targeting as a conspicuous phenotypic reporter of active silencing.

**Figure 1.**
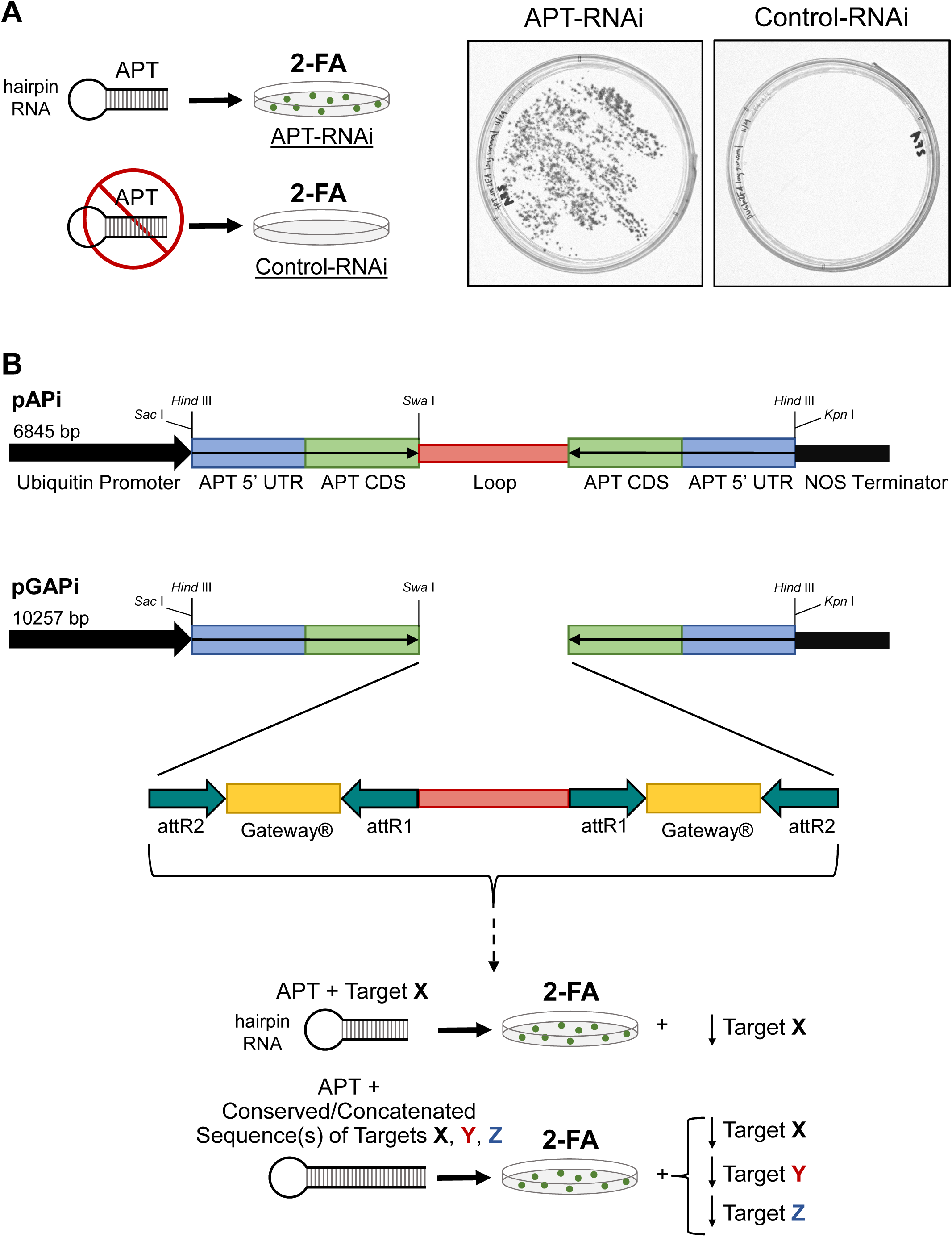
Proof-of-principle and construction of APT-based RNAi (APTi) vectors to enable positive selection of actively silencing plants. A, Illustration of the APT-interference positive selection principle: plants are transformed with a vector that creates a long dsRNA hairpin targeting the APT gene, thereby reducing endogenous APT levels and subsequent production of cytotoxic nucleotides when supplemented with 2-fluoroadenine (2-FA). Targeting APT using RNAi is sufficient for transformed plants to grow on standard PpNH_4_ medium supplemented with 1.25 μg/mL 2-FA. B, Schematics of the APT-based RNAi vectors. The pAPi (<u>p</u>lasmid <u>AP</u>T RNAi) and pGAPi (<u>p</u>lasmid <u>G</u>ateway® <u>AP</u>T RNAi) constructs were created using the pUGi and pUGGi vectors, respectively, from Bezanilla et al., 2005 as templates. The thin black arrows indicate the direction of the open reading frame, and the inverted repeat regions of both constructs are flanked by a constitutive maize ubiquitin promoter (thick black arrow) and a NOS terminator sequence (black rectangle). Both constructs target the 5’ UTR (blue rectangle, 179 bp) and coding sequence (green rectangle, 210 bp) of the APT gene. pAPi contains only the loop (red rectangle, 402 bp in pAPi, 392 bp in pGAPi) region within the inverted repeat, whereas pGAPi contains inverted Gateway® sites to facilitate insertion of target sequence. The target may be unique to gene X, or if the target sequence is conserved the user can simultaneously silence multiple targets in tandem with APT silencing, thereby enabling survival of silencing plants.

Our previous APT silencing experiment demonstrated feasibility, but further development of the technique was constrained by the available construct. As mentioned above, the first iteration of the APT-based interference (APTi) was inserted into a vector created specifically for RNAi (Bezanilla et al., 2005). This construct included a pair of inverted Gateway® sites coupled to a target sequence (GUS) for an internal reporter of active silencing. The reporter sequence targeted a nuclear-localized GFP:GUS fusion protein, thereby requiring the use of a special transgenic line for RNAi experiments (Bezanilla et al., 2003). In principle, our novel APTi would not require any specific moss line, and instead would permit the researcher to perform RNAi experiments in any genetic background. Therefore, we replaced the GUS reporter target sequence with the APT target sequence, while maintaining the inverted Gateway® cassettes and loop region and named the construct pGAPi (Fig. 1B). This construct allows straightforward insertion of any gene sequence and ensures fusion to the APT target, thus permitting inference of gene silencing within surviving plants. Importantly, entire gene families can be targeted by simple insertion of a conserved sequence of approximately 400 bp, or concatenation of individual target sequences followed by cloning into our pGAPi (Fig. 1B). Additionally, we created another APTi construct, named pAPi, which lacks the internal Gateway® cassettes to serve as a “positive control” for any RNAi experiment (Fig. 1B). Together, these constructs serve as the foundation for our novel APT-based RNAi (APTi) that advances positive selection as an effective reporter of gene silencing.

### APTi Experimental Design for High-throughput Phenotyping

Although our APTi system clearly selects for silencing plants cultured over a two-week period by visual inspection (Fig. 1A), we sought to establish a rapid, semi-automated microscopy assay using APTi for plant phenotyping. A fundamental attribute of any high-throughput assay is the effective and automated discrimination between objects of interest and background that co-occupy the same space. Our APTi system simplifies the problem of automated separation. Unlike fluorescence reporter systems where the reporter signal is continuous and exhibits natural variation, APTi reduces separation of silencing and non-silencing plants to the binary decision of alive or dead. We reasoned that chlorophyll autofluorescence could function as a proxy for plant survival, thereby enabling automated detection of alive, and therefore actively silencing, plants. In principle, all plants successfully transformed with a construct silencing APT, pAPi (Fig. 1B), will survive on 2-FA medium, whereas plants transformed with an RNAi construct silencing a non-existent reporter, pUGi (Bezanilla et al., 2005), will die.

We tested this by using the experimental design illustrated in Fig. 2. *P. patens* protoplasts were transformed with either pAPi or pUGi, allowed to regenerate for 4 days, then transferred to growth medium supplemented with 1.25 μg/mL 2-FA. Importantly, during the optimization phase of this assay we observed substantial variability in the outcome of 2-FA selection. We empirically determined that starting the selection four days post-transformation and making the 2-FA selection plates fresh on the day of selection mitigated essentially all experimental variability. We strongly suggest first optimizing the 2-FA selection concentration when applying the APTi system, as our results were all obtained using a single lot of 2-FA from Oakwood Chemical, SC, USA.

**Figure 2.**
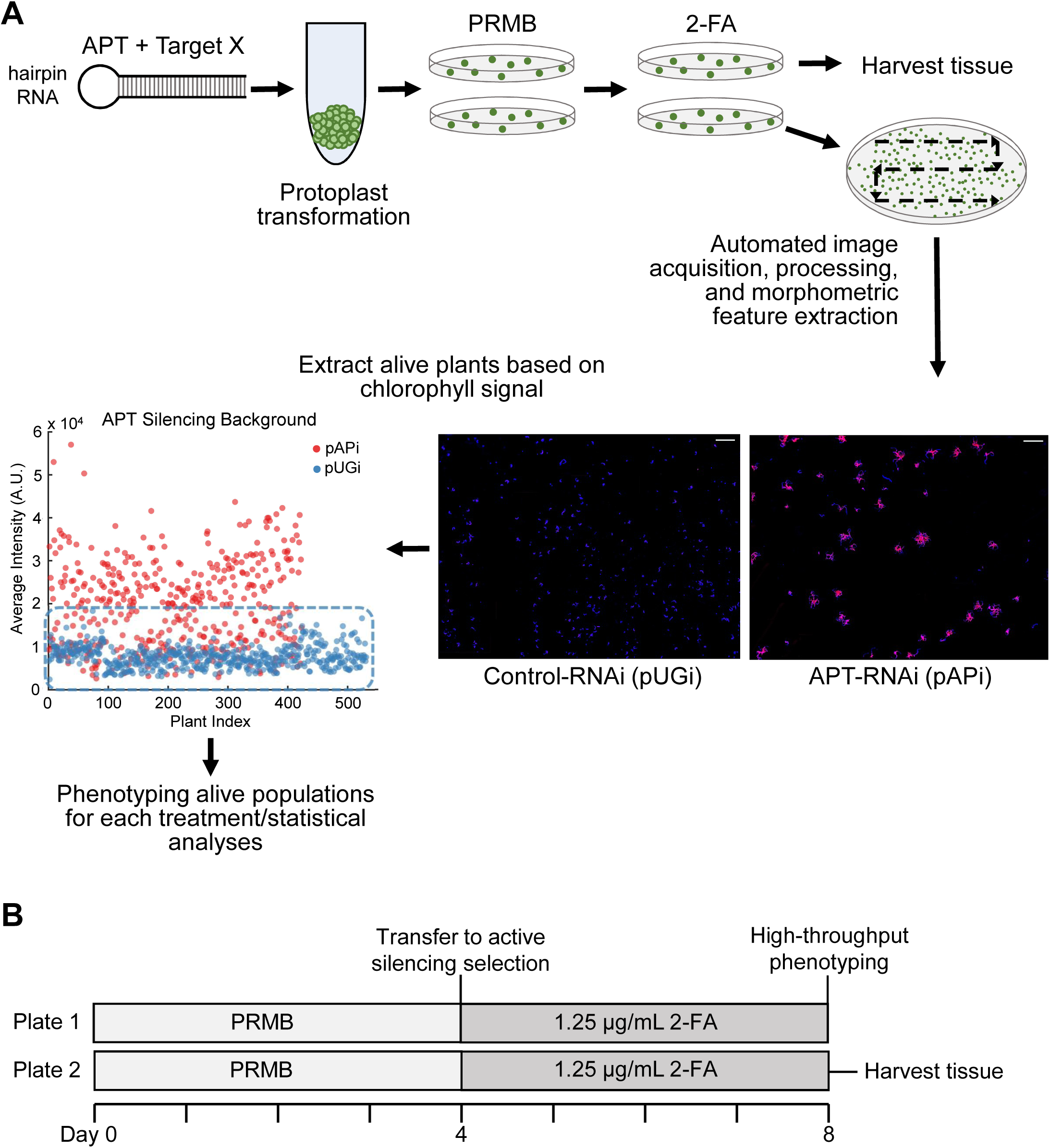
Experimental strategy and timeline for the novel APTi system. A, Optimized experimental pipeline for transient transformation of *P. patens* protoplasts and high-throughput acquisition and analysis for the APTi system. Images are a subsection of the entire plate area and demonstrate the characteristic difference in survival between conditions that do (pAPi) or do not (pUGi) silence APT. The scatter plot determines the background threshold value, which is based upon the maximal observed chlorophyll signal in the Control-RNAi condition (pUGi, blue dots). Each point corresponds to an individual plant, where the points are pooled from three independent experiments. B, Timeline of events for an APTi assay. Typically, each condition is plated in at least duplicate for adequate sample abundance for both imaging and molecular analyses.

Following four days on 2-FA supplemented growth medium, cultures were removed from the growth chamber for protein extraction and microscopy analysis (Fig. 2B). To facilitate high-throughput phenotyping and remove human bias, we used an epifluorescent microscope equipped with an automated stage integrated with image tiling and stitching software. This enabled large region-of-interest acquisition, with the size of the ROI only constrained by the memory available to the computer. For our experiments, every composite image is constructed from a 12×12 grid of single images, with a 15% overlap, which corresponds to an ROI surface area of approximately 1.8 cm^2^.

To simplify downstream image segmentation, prior to imaging all plants were stained with calcofluor to label the cell wall. Each individual image contained two channels, the chlorophyll autofluorescence and the calcofluor signal. Visualization of the chlorophyll signals of APT-RNAi and Control-RNAi clearly reveals the efficacy of the APTi system: not a single plant with endogenous levels of APT survives and does not markedly grow beyond the initial regeneration size (Fig. 2A). This result is reproducible, as the chlorophyll signal across three independent experiments for Control-RNAi plants consistently clustered below a characteristic intensity (Fig. 2A). We used chlorophyll intensity parameter as a threshold, from which we functionally partitioned a mixture of plants on a plate into alive (silencing) and dead (non-silencing) plants (Fig. 2A). Classification as alive or dead ensured further image analysis of morphometric parameters or other features was confined to only alive, and therefore silencing, plants. Therefore, it stands to reason that when using our APTi system any statistically supported observed difference between plants transformed with APTi + gene target and plants transformed with pAPi is directly attributable to silencing of the targeted gene.

### The APTi System Effectively Isolates and Enables Rescuing of a Polarized Growth Mutant

We demonstrated that silencing of APT permits potent positive selection of actively silencing plants (Fig. 2A), but we sought to extend APTi to mutant analysis. We reasoned that insertion of a target sequence into our APTi vector (Fig. 1B) would produce in tandem silencing of APT and the target gene. To test this hypothesis we exploited the well-characterized *myosin XI* mutant that produces a dramatic loss of polarized growth phenotype (Vidali et al., 2010). Furthermore, the *myosin XI* mutant was previously generated using a fluorescent reporter-based RNAi strategy, allowing a direct comparison of methodology.

Insertion of a concatenated 5’ UTR sequence derived from both isoforms of myosin XI into our APTi vector pGAPi (“myoUTi”) resulted in a striking recapitulation of the *myosin XI* phenotype (Fig 3A). Impressively, nearly every surviving plant manifested the characteristic “bunch of grapes” morphological phenotype (Fig. S1). This is exemplified by the relatively narrow distributions of the APTi myosin XI knockdown in the two morphology parameters, solidity and area (Fig. 3B). We speculate the survival advantage inherent to our APTi method could reduce phenotypic variability sometimes observed in RNAi experiments.

**Figure 3.**
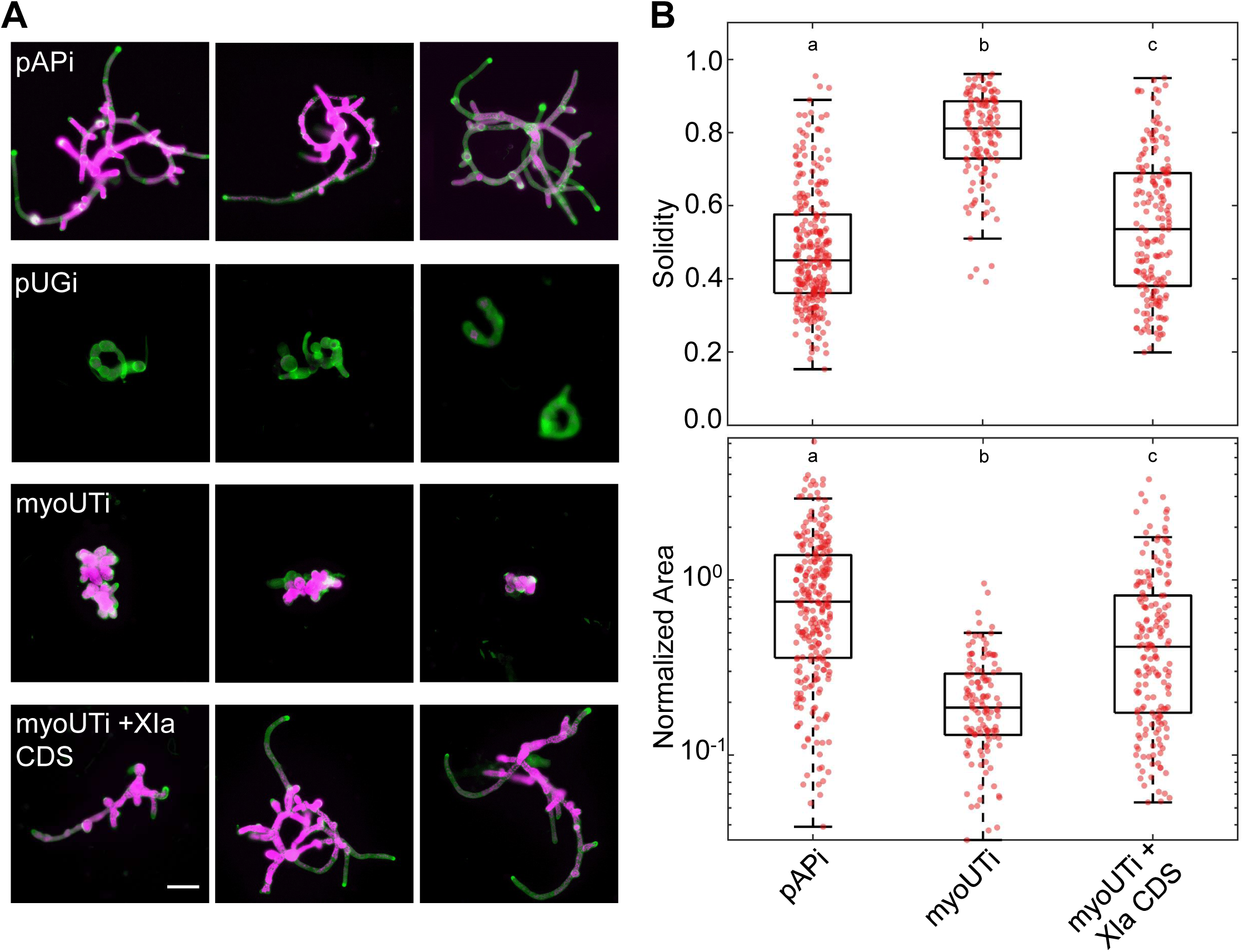
The APTi system robustly captures and rescues the strong loss-of-growth and polarization mutant of myosin XI in *Physcomitrella patens*. A, Representative images of 8-day old plants regenerated from protoplasts transformed with APTi vectors targeting APT alone (pAPi), APT in tandem with a concatenated 5’ UTR sequence for myosin XIa and XIb (myoUTi), targeting a non-existent GUS sequence (pUGi), or myoUTi co-transformed with a construct overexpressing myosin XIa’s coding sequence. All images are cropped from composite images that capture a large sample area for an individual condition, as shown in Figure 2A. For every condition, each image corresponds to an independent experiment. Chlorophyll autofluorescence is colored magenta and calcofluor signal is colored green. Scale bar = 100 μm. B, Quantification of morphometric parameters solidity and area from three independent experiments. Area is normalized to the mean area of the pAPi condition, which represents near wild-type growth morphology. pAPi, n= 270; myoUTi, n = 125; myoUTi + XIa CDS, n = 178. Letters indicate statistical difference (P < 0.001) between groups as determined by one-way ANOVA with post-hoc Tukey test.

We inquired whether this phenotype could be rescued in the APTi system by co-transforming the “myoUTi” construct with a plasmid expressing only the coding sequence of myosin XIa, “XIa CDS.” We observed near complete rescue of the *myosin XI* phenotype, demonstrating an absence of off-target effects and, more importantly, highlighting the rapid phenotyping utility of the APTi system when coupled with *P. patens*. Within 8-days our APTi system isolated through positive selection a relatively homogenous population of mutant plants, which are amenable to rescue. Our APTi system produced average myosin XI mutant morphological parameters similar to a myosin XI mutant derived from RNAi using an internal GFP reporter system (Vidali et al., 2010), but without non-silencing background plants. Together, these results support our prediction that aberrant morphologies observed using the APTi system are caused by silencing of the conjugated target, in our case myosin XI.

### Survival of Plants Using APTi Directly Corresponds to Potent Reduction of Target Protein

A significant limitation of all reporter systems to date, such as the popular GFP reporter, is the presence of a substantial background population of plants not undergoing RNAi. This places a substantial burden on the experimenter, as it requires individual, microscopic isolation of plants that pass some reporter threshold for active silencing. Characterization of strong growth mutants, such as myosin XI and profilin, only compound the problem because not enough plant tissue can be isolated for reliable protein quantification (Vidali et al., 2007; Augustine et al., 2008; Vidali et al., 2009; Vidali et al., 2010). Instead, labor intensive immunostaining or indirect estimation of protein translation silencing via transcript abundance is required. As our novel APTi approach removes essentially all background, we asked if the previously intractable problem of reliable protein quantification could be trivialized to harvesting all material present on the 2-FA plate. If successful, we predicted a substantial reduction in total myosin XI protein when treated with APTi, which explains the observed mutant morphology (Fig. 3).

To accurately quantify myosin XI protein abundance using our APTi method, we first established the linear range of our antibodies. To reflect our RNAi experimental conditions, we transformed wild-type moss with pAPi, grew on medium supplemented with 2-FA, then harvested the entire plate at 8-days post-transformation. Total protein was determined using a Bradford microplate microassay (Bio-Rad), then a range of total protein (1-20 μg) was probed using both an in-house developed antibody against myosin XIa’s coiled-coil tail (CCT) region and a publicly available anti-alpha tubulin antibody (DSHB: AA4.3). This approach revealed an approximate linear range for both antibodies from 5-20 μg total protein (Fig. S2A). Importantly, under equivalent conditions our limit of detection for endogenous, wild-type myosin XI is approximately 1 μg total protein (Fig. S2A). This step is essential, as it allows for confident, semi-quantitative estimation of the extent of protein reduction. Without this, the wild-type protein of interest could be loaded at or very near the limit of detection, resulting in a complete absence of protein signal in the RNAi condition even if the true reduction is modest.

We performed an APTi experiment for myosin XI, as shown in Fig. 3, and harvested at 8-days post-transformation. Every condition was implemented in at least duplicate to allow for both harvesting of plants and our phenotyping assay (Fig. 2). In this way, the results of our protein analysis directly reflect the internal protein abundance and corresponding morphologies we observe in the phenotyping assay (Fig. 3). We observed a dramatic decrease of myosin XI in the “myoUTi” condition, which when normalized to α-tubulin results in a maximal 93% reduction relative to the control (“pAPi”). Additionally, the rescued mutant morphology precisely corresponds to an almost wild-type restoration of myosin XI levels (Fig. 4). Of note, these results are consistent across independent experiments and results in an average silencing efficiency of 90% (Fig. 4). We explored the longevity target silencing using APTi by probing myosin XI at two weeks post-transformation. Although less potent than at 8-days post-transformation (Fig. 4), myosin XI levels are still substantially reduced (Fig. S2B,C), opening the possibility for long-term phenotyping. At the longer time, the rescue condition is noticeably weaker than at the short time point. We attribute this to loss of the myosin XI expression plasmid, as it is not under selection. Taken together, these data establish that aberrant phenotypes observed using our APTi system are directly attributable to reduction of target protein abundance.

**Figure 4.**
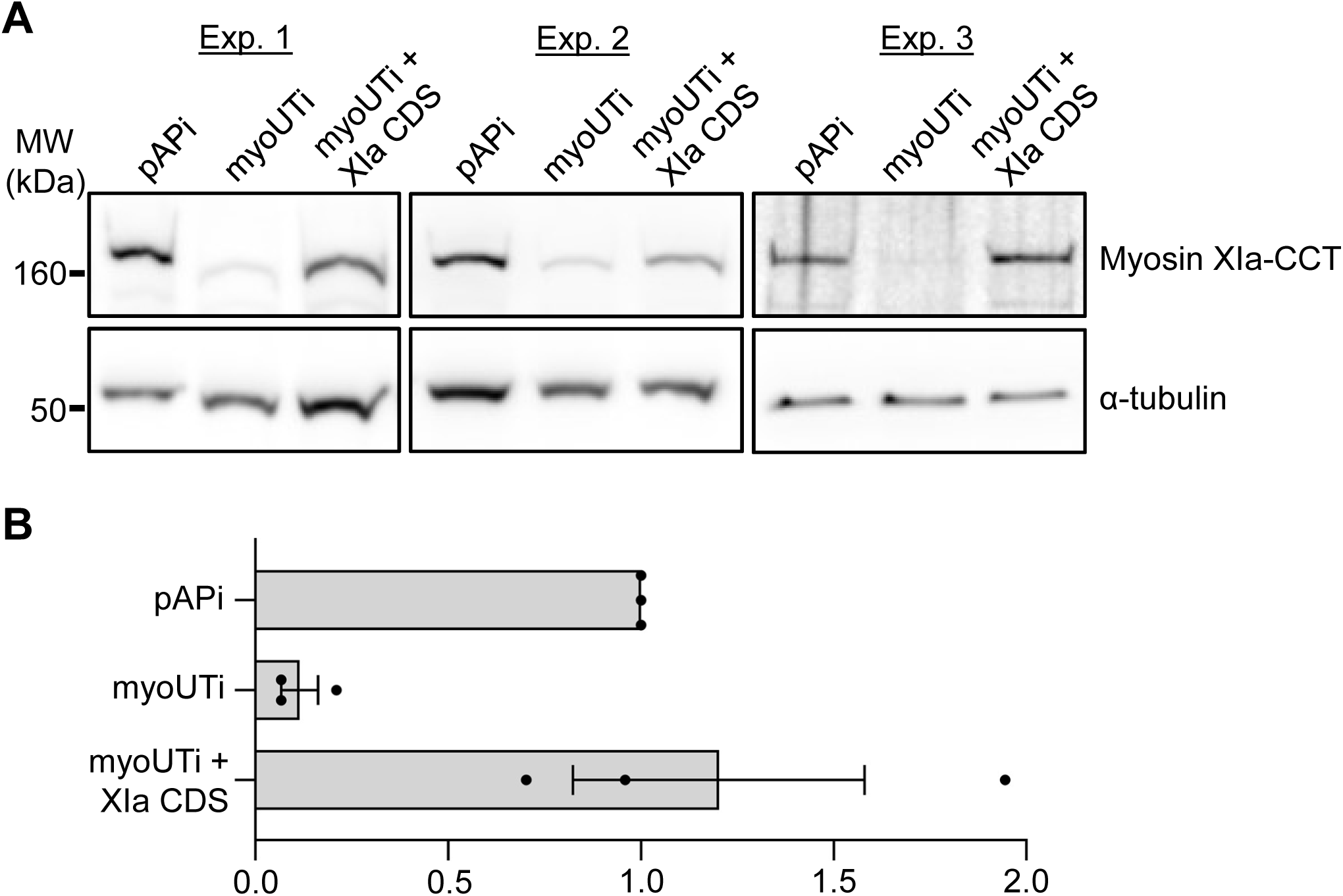
Plant survival using the APTi system directly corresponds to reduction in target protein abundance and is rescuable. A, Western blots demonstrating the reduction and restoration of myosin XI protein levels when using the novel APTi system. Each experiment represents an independent transformation and subsequent plant harvesting and western blotting. 10 μg of total protein was loaded per lane. Myosin XI was probed using a polyclonal antibody generated against the myosin XIa coiled-coil tail (CCT) fragment from *Physcomitrella patens* and α-tubulin was used as a loading control. B, Densitometry of western blot signals was performed using ImageJ and shows an approximate 90% reduction of myosin XI and a complete rescue of myosin XI protein levels when compared to the pAPi control.

## Discussion

RNAi offers an invaluable complement to traditional gene knockout studies. However, substantive advancements in RNAi methods are trailing the explosion of CRISPR/Cas-based technologies. Here we established the first survival-based RNAi methodology that robustly selects for actively silencing plants. We accomplished this by engineering vectors that elicit a pro-survival response when processed by the organism’s endogenous RNAi machinery. Using the previously characterized *myosin XI* mutant in *P. patens* as a case study, we showed that in tandem fusion of a myosin XI target with the pro-survival sequence resulted in potent selection of morphologically mutant plants. Furthermore, we demonstrated that surviving plants actively targeting myosin XI through our novel vectors contained approximately 7% of normal myosin XI protein abundance. This technology represents a dramatic improvement in silencing efficacy and experimental implementation over previous RNAi methods.

RNAi has been extensively employed in the model moss *Physcomitrella patens* for both discovery and validation of gene function (Vidali et al., 2007; Augustine et al., 2008; Prigge et al., 2010; Vidali et al., 2010; Augustine et al., 2011; Wu et al., 2011; Miki et al., 2015; Bascom et al., 2019), to cite a few examples. We attribute the popularity of dsRNA-based RNAi in *P. patens* to a method that uses an internal fluorescent reporter of RNAi to obtain results in one week (Bezanilla et al., 2003). We sought to fundamentally improve upon RNAi reporters by creating a reporter where survival itself is indicative of active silencing. We demonstrated the feasibility and effectiveness of this approach by silencing the APT gene in the presence of 2-FA in the medium. Furthermore, we engineered two plasmids that facilitate insertion of any target of interest in tandem with the APT-silencing sequence. We call this novel approach APT-based RNA interference, or APTi. Both APTi plasmids and the myosin XI-RNAi plasmid are publicly available from the plasmid repository Addgene.

We obtained potent positive selection of actively silencing plants in *Physcomitrella patens* by engineering RNAi vectors that exploit the function of the APT gene. However, the physiological consequences of APT silencing remain in question. Although a salvage enzyme, APT’s function presents a more energetically efficient means of nucleotide production than *de novo* synthesis (Ashihara et al., 2018). Consequently, knockouts of APT in vascular plants demonstrate severe defects in pollen germination and pollen tube growth, presumably a result of impairing the energy-intensive fast growth of the pollen tube (Moffatt and Somerville, 1988; Zhou et al., 2006). Interestingly, an alternative mutant allele of APT that results in partial reduction of APT activity imparts enhanced growth and stress tolerance (Sukrong et al., 2012). We suspect this hypomorphic allele better represents the internal state of an APT-silenced plant. Taken together with our observations of APT-silencing plants reproducing results achieved with an orthogonal RNAi system, we conclude the reduction of APT results in no clear physiological defects within the scope of our assay.

We expect our APTi strategy to be applicable to other organisms given the ubiquity of the APT gene (Schaff, 1994). Like other organisms, such as in humans, *Physcomitrella patens* contains only one copy of APT making it especially amenable to the APTi strategy as we showed. Interestingly, the vascular plant *Arabidopsis thaliana* contains five APT genes (Allen et al., 2002). We suspect APTi could be applied in *A. thaliana* by constructing an APT-silencing module comprised of concatenated sequences targeting specific isoforms. Additional work is necessary to translate to other systems, but we submit that our efforts establishing APTi in *P. patens* will greatly benefit the community in understanding fundamental and conserved biological processes (Orr et al., 2020).

Previous work using the fluorescent reporter-based RNAi identified silencing plants based on loss of fluorescence (Bezanilla et al., 2003). However, slight reduction in fluorescence confounded interpretation because it could be attributed to natural variation in the reporter signal or reflect a real, but modest, silencing effect. Furthermore, in our hands we observed spontaneous loss of reporter signal in the moss reporter line over long periods of continuous propagation. Without careful observation and subcloning to remove chimeric reporter cultures, a researcher could inadvertently conclude silencing when none is occurring. Our APTi system simultaneously removes the requirement of a dedicated moss reporter line and dismisses any ambiguity inherent to a continuous reporter signal. Additionally, previous RNAi methods can result in background plants that survive antibiotic selection for the plasmid containing the RNAi transgene but do not silence the reporter. This is likely a result of transcriptional gene silencing, whereby the plant rescues itself from RNAi transgene expression but expression of the independently regulated antibiotic resistance is unmodified (Morel et al., 2000; Fusaro et al., 2006; Small, 2007). We hypothesize our method enhances silencing efficiency by engineering a fitness punishment for the organism to silence the RNAi transgene. To this end, with APTi survival on 2-FA is directly coupled to expression of the APT-RNAi transgene. Therefore, the organism cannot survive if it silences the expression of the RNAi cassette, thus ensuring expression of the APT + gene target hairpin. Although we did not determine the extent of APT silencing, we know the reduction is sufficient to promote survival on 2-FA and produce a 90% reduction of the fused target, myosin XI.

We speculated that the enhanced fitness benefit imposed by our APTi system will result in more consistent and potent silencing efficiencies of target genes. Consistent with this, we observed a homogenous population of myosin XI mutants and strong, 90% reduction of endogenous myosin XI. APTi offers a noticeably higher silencing efficacy when compared to previous reports using dsRNA and GFP-based reporters in *P. patens* (Vidali et al., 2007; Vidali et al., 2010; Nakaoka et al., 2012). Importantly, previous reports could only estimate protein silencing based on a small subset of individual plants deliberately chosen by the experimenter, which may not accurately reflect the average silencing effect (Vidali et al., 2007; Augustine et al., 2008; Vidali et al., 2009; Vidali et al., 2010). This microscope-based methodology was required because mutants resulting in small morphologies failed to yield adequate plant material for immunoblots, and were surrounded by non-silencing background plants. We demonstrated that APTi’s positive selection enables simple harvesting of the entire plate that can be easily scaled for reproducible protein quantitation. Based on our observed myosin XI knockdown, our APTi silencing efficacy is comparable to the most effective amiRNAs (Zhang et al., 2018), but without the need for prior engineering and screening of multiple amiRNAs.

We showed that the APTi system is well suited for high-throughput phenotyping. We fully anticipate this area to be iteratively improved, not just with respect to the volume of acquisition but with increased computational sophistication. The large obtainable data sets are ripe for both classic exploratory data methods and cutting-edge deep learning techniques. For example, we analyzed living plants by first segmenting the images by traditional thresholding. We then filtered and classified the hundreds of plants based on a characteristic biological feature, chlorophyll autofluorescence, which we derived from the Control-RNAi dying population. Although less intuitive, deep-learning offers the possibility of automating image segmentation and classification, perhaps resulting in discovery of novel mutant features (Moen et al., 2019).

## Conclusions

This work represents a fundamental transition from visual screening for RNAi plants to positive selection of actively silencing plants. We achieved this by engineering vectors that produce hairpin RNA targeting the APT gene fused to any gene target(s) of interest. This results in effective isolation of all surviving plants undergoing RNAi of the target gene when grown in the presence of 2-FA. Importantly, the efficacy of gene-silencing was consistently greater, maximal 93% reduction of target protein, with the APTi system than previous reports silencing the same gene using a fluorescent screening method. We believe our APTi system provides a flexible, fast, and effective platform with unprecedented low background and variability to facilitate high-throughput characterization for loss-of-function mutants.

## Materials and Methods

### Plant materials and culture conditions

Two *Physcomitrella patens* lines were used in this study: (1) NLS4 (Bezanilla et al., 2003) in Figure 1A; (2) “wild-type” Gransden (Ashton et al., 1979) all other experiments. All lines were cultured as previously described (Vidali et al., 2007). In brief, tissue was propagated weekly by homogenization and transferred to solid PpNH_4_ medium overlaid with cellophane. Cultures were grown at 25°C under long-day light (90 μmol m^-2^ s ^-1^) conditions (16 h : 8 h, light : dark).

### APT-based RNAi construct design

The APT transcript fragment (Phytozome: Pp3c8_16590) containing 5’UTR and coding sequence (CDS) was amplified by PCR with forward (APT_BSK_F) and reverse (APT_BSK_R) primers and cloned into pBlueScript K+ using restriction enzymes *SacI/EcoRV* (generating the APT pBSK+ plasmid). The pUGGi Gateway cassette and loop domain lacking the GUS regions was amplified in two pieces using PCR with primer sets Gateway1_F/R and Gateway2_F /R (both reactions at 60°C, 2min elongation), and both fragments were ligated individually into pBlueScript K+ (generating the Gateway F pBSK+ and Gateway R pBSK+ plasmids). The pUGGi loop was amplified by PCR with primers Loop_F/R and inserted into pBlueScript K+ (generating the Loop pBSK+ plasmid). The APT and Loop pBSK+ constructs were transformed into DH5α *E. coli*, whereas both Gateway pBSK constructs were transformed into ccdB *E. coli*. All constructs were blue-screened for successful clones by plating transformants on LB+Carb+Chlor plates and adding 40µL each of IPTG and X-Gal. Next, using the *SacI/XhoI* restriction sites the Gateway F fragment was inserted into the Gateway R pBSK+ backbone, generating the complete Gateway pBSK+ plasmid. Importantly, this construct contains the entire loop region derived from pUGGi, as the two intermediate Gateway constructs discussed above each contain half of the loop region. It was necessary to isolate the pUGGi Loop in its own plasmid because we wished to create an additional control plasmid lacking the Gateway cassettes.

The first APT fragment was then excised from APT pBSK+ using *HindIII/SwaI*, then transferred into the Gateway pBSK+ plasmid, cut with *HindIII/PmeI* (generating Gateway-1APT pBSK+). Next, the second APT fragment was excised from APT pBSK+ and transferred into the Gateway-1APT pBSK+ plasmid using *SacI/SwaI* sites (generating Gateway-2APT pBSK+). The Gateway-2APT fragment was digested then ligated into the pUGGi backbone using *SacI/KpnI* sites, generating the pGAPi plasmid. To generate our positive control plasmid without Gateway sites, two APT fragments were inserted into the Loop pBSK+ plasmid using the same procedure as described above—first by *HindIII/SwaI* and *HindIII/PmeI* sites, then by *SacI/SwaI* sites - producing the Loop-2APT pBSK+ plasmid. Then the Loop-2APT fragment was cloned into the pUGGi backbone using *SacI/KpnI* sites, generating the pAPi plasmid. At all points in this process where new plasmids were made by restriction-based cloning, those plasmids were confirmed by restriction and sequence analysis. Both pAPi and pGAPi are publicly available from Addgene (pGAPi #127547; pAPi #127548; myoUTi:pGAPi #127549) and sequences are also available at Genbank (pGAPi = MK975250; pAPi = MK975251; myoUTi:pGAPi = MK975252).

### APTi phenotyping assay

One week-old moss was protoplasted and transformed as described (Liu and Vidali, 2011), with transformed protoplasts being plated at 1.4 x 10^5^ protoplasts per 100 mm petri dish of PRMB in liquid plating medium. Each condition was plated in at least duplicate, with the myoUTi condition being plated in triplicate to allow enough plant material of the mutant plants to be harvested for immunoblotting and imaging. Four days post-transformation the cellophanes of each plate were transferred to growth medium (PpNH_4_) supplemented with 1.25 μg/mL 2-fluoradenine from a 5 mg/mL in DMSO stock (Oakwood Chemical). As mentioned previously, we emphasize that each lab should first optimize the 2-FA selection concentration, as our results were all obtained using a single lot of 2-FA from Oakwood Chemical, SC, USA.

Eight days post-transformation plants were stained with 10 μg/mL calcofluor from a 1 mg/mL in H_2_0 stock (Fluorescent Brightener 28, Sigma) and imaged with a 10X A-Plan (0.25 NA) objective of an epifluorescent microscope (Zeiss Axiovert 200M) coupled to a CCD camera (Zeiss AxioCam MRm). This microscope was equipped with an automated stage and integrated with the AxioVision software through the MosaicX module, enabling precise acquisition, tiling, and stitching of individual images to create a large composite. Our composite images contained 12×12 individual images, acquired with a 15% overlap. Each individual image consisted of two channels, the calcofluor and chlorophyll signals. The chlorophyll channel was acquired with a 480/40 bandpass excitation, a 505 longpass dichroic mirror, and a 510 long-pass emission filter cube and with a fixed 150 ms exposure for all experiments. The calcofluor signal was acquired with a standard DAPI filter and automatically adjusted for each experiment to maximize contrast. Stitched images were first processed and segmented using a custom ImageJ macro (available upon request). Our macro only discarded segmented objects approximately the size of two adjoining protoplasts or smaller, as to not bias against the discovery of small mutants and remove non-surviving protoplasts. Each segmented image was visually inspected for overlapping or truncated plants, and if present they were discarded from further analysis. Subsequent filtering of alive plants and visualization of the data was performed using MATLAB (MathWorks). Statistical testing (one way ANOVA-Tukey) was performed with GraphPad Prism.

### Analysis of myosin XI protein abundance

Eight days post-transformation, all tissue was harvested by scraping, flash-frozen in liquid nitrogen, and stored at -80°C. To create protein extracts, the frozen tissue was ground to a powder in liquid nitrogen, then resuspended in extraction buffer (250mM Sucrose, 20mM EGTA, 50mM PIPES, 150mM NaCl, 60mM MgCl2, and 1% w/v Casein) supplemented with fresh DTT (2mM final) and protease inhibitors. The powder from the pAPi condition was resuspended in 300 μl, whereas the myoUTi and rescue conditions were always resuspended in 200 μl extraction buffer. Extracts were vortexed for 15 seconds, then placed on ice for 30 seconds, with this repeated twice more. Extracts were then spun at 13,000 RPM for 10 mins at 4°C, followed by removal of 175 μl of the clarified supernatant; 120 μl of extract was immediately combined with SDS loading buffer and boiled, then stored at -80°C. The remaining extract was used for total protein determination using the Bradford microplate microassay procedure (Bio-Rad).

To probe myosin XI protein directly, we developed an antibody against a 6xHis fusion of the *P. patens* myosin XIa-CCT (Capralogics, Inc. Hardiwick, MA). We used an antibody against alpha tubulin as our loading control (DSHB: AA4.3). We determined the approximate linear range of both antibodies by first loading 1-20 μg of total protein from a pAPi treated moss extract on a 4-12% Bis-Tris SDS-PAGE gel (ThermoFisher). Protein was then transferred to nitrocellulose overnight at 4°C, followed by blocking with 5% milk in TBS-T at room temperature for 1hr. The nitrocellulose was cut at the 80 kDa marker, with the higher molecular weight piece incubated with myosin XIa-CCT primary antibody (1:10,000) and the α-tubulin (0.5 μg/ml final) incubated with the lower molecular weight fragment for 1hr at room temperature. Following primary incubation, blots were washed three times in TBS-T, incubated in secondary antibody (goat anti-rabbit for XIa-CCT, goat anti-mouse for α-tubulin) at 1:100,000 dilution for 1hr at room temperature, then washed a final three times. Blots were developed using homemade ECL reagent and chemiluminescent images were acquired using an Azure 600 (Azure Biosystems). Densitometry was performed using ImageJ (Schneider et al., 2012) to allow comparison of relative protein abundance.

## Supplemental Data

**Supplemental Figure S1**. The APTi system reduces variability in observed mutant phenotype.

**Supplemental Figure S2**. Determination of the linear range of used antibodies and long-term reproducibility of western blots using the APTi system.

**Supplemental Table S1**. List of primers used for vector construction and sequence verification.

**Table 1.**
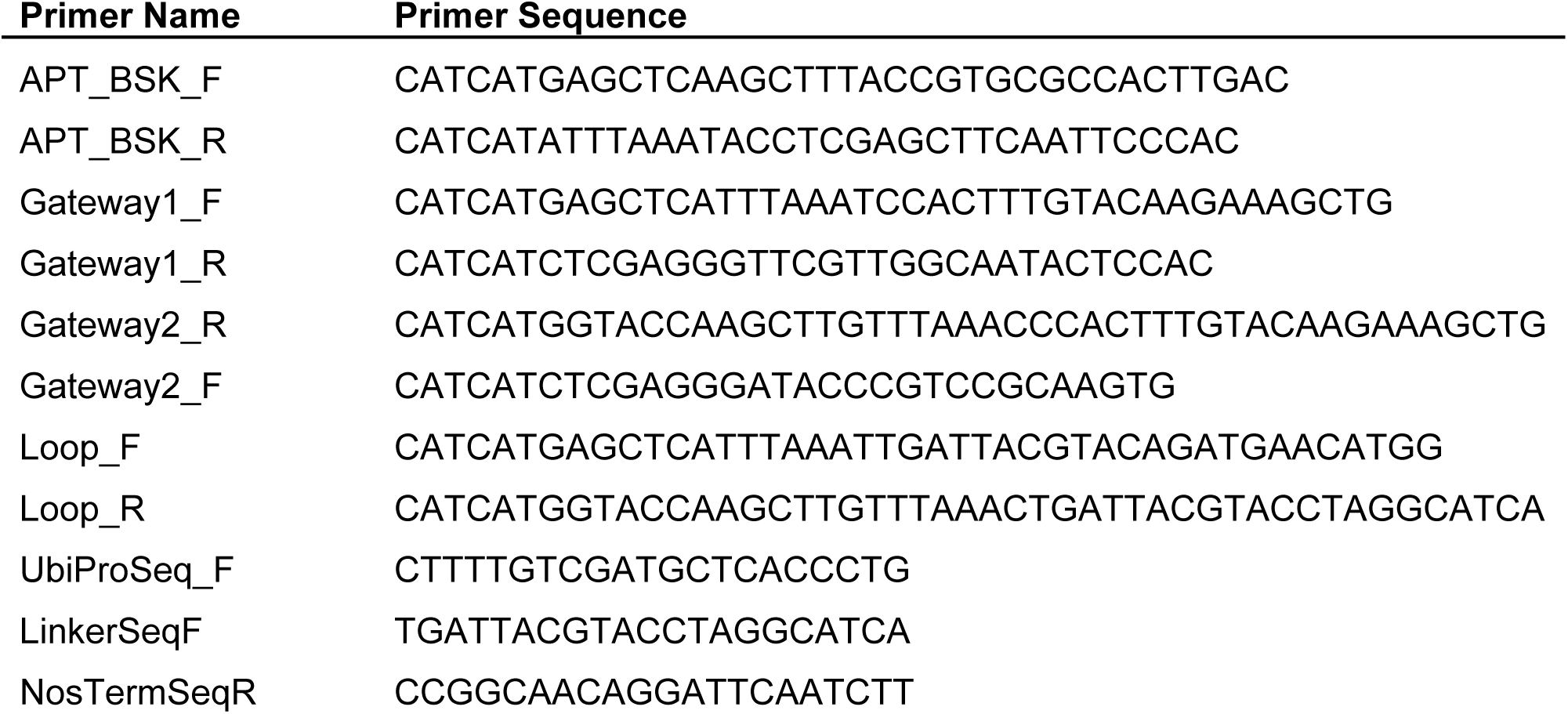
List of primers used for vector construction and sequence verification.

## Acknowledgements

We thank the members of Luis Vidali’s group at Worcester Polytechnic Institute and Mary Munson’s group at University of Massachusetts Medical School for helpful discussions. We also thank Victoria Huntress for her management of the Microscopy core facility at Worcester Polytechnic Institute.

